# Rehoming laboratory rats: Exploring perceptions of rehomers, animal technicians and biomedical researchers

**DOI:** 10.1101/2025.07.31.667926

**Authors:** Georgia Greenan, Caroline Quigley, Ignacio Vinuela-Fernandez, John Menzies

## Abstract

Laboratory animals can be rehomed at the end of a study. However, there is relatively little research on stakeholders’ perceptions of rehoming. We explored the views of three key groups: the people who volunteer to rehome these animals, the animal technicians who care for these animals prior to rehoming, and the researchers who are normally responsible for most of the interactions between people and these animals. To do this, we carried out a thematic analysis of semi-structured interviews. Our aim was to obtain insights that could inform reflection and guide future research priorities around rehoming programmes. Our study demonstrated support for laboratory rodent rehoming in all stakeholder groups. Rehomers’ and technicians’ comments focused chiefly on the potential benefits of rehoming to the animal and benefits to institutional openness. Researchers made similar comments, but in this group, these comments were tempered by concerns around the welfare of the animals after rehoming, the fate of animals that cannot be rehomed, rehomers’ health and well-being, and the potential risks associated with the transparency of individual’s and the institution’s use of animals in research. We discuss these themes in the context of the ethics of laboratory animal use, manifesting a Culture of Care, and challenges around enhancing institutional openness on animal research.

## Introduction

Humans work with animals in many different ways. In some contexts, animals are allowed to ‘retire’ and be rehomed in sanctuaries or homes when they are no longer able or needed to work [1–3]. Rehoming is normally perceived to benefit the animal being rehomed, and to benefit the people who worked with, and rehomed, the animal [4–6]. Accordingly, rehoming is usually framed as an ethically positive action that is compatible with ideas around fairness and flourishing for animals as moral patients [7–9].

Mice and rats accounted for 77% of the total number of animals used in 2.68 million procedures in 2023 in the UK [10]. It is likely that almost all of these animals were euthanised as part of or after these procedures, often in order to collect biological material required for the study. However, some would have been killed simply because their scientific usefulness had come to an end or because they fell into the ‘bred, but not used’ category, regardless of whether they were bred but could not be used, or bred but were not needed [11,12]. However, the legal and ethical frameworks that regulate the scientific use of animals do not demand euthanasia at the end of a study. In addition to ‘re-use’ as an alternative to death, rehoming is specified as potential outcome in UK and EU regulations [13,14]. Quite strikingly, Tess Skidmore points out that under the definition of rehoming in UK legislation, “farm animals used in research may be ‘rehomed’ to a slaughterhouse, or animals may be ‘rehomed’ to another research facility abroad to undergo experimental re-use” [8], but in the context of our study, we are referring to rehoming as transferring an animal to a private home to be cared for as a pet. A laboratory animal is a candidate for rehoming if it is healthy, adequately socialised and it poses no danger to public health, animal health or the environment [15]. Importantly, UK and EU regulations rightly require that measures must be in place to protect the wellbeing of the animal once rehomed.

A recent survey showed that rehoming programmes are fairly common in many UK animal research facilities[16]. However, the number of laboratory animals rehomed was quite small – just 2,322 in 2015-2017. Notably, species which are commonly perceived as pets are rehomed at the highest proportion (Skidmore discusses reasons for this in [17]). For example, from 2015 to 2017, 171 cats (38% of the total number kept nationally for research) were rehomed. In contrast, the forty-one animal facilities included in the study reported the rehoming of just 103 rats and 22 mice in the same period, representing just 0.008% and 0.0003% of the total number kept in the UK for research in that period. This equates to an average of <1 rat rehomed per facility per year. A 2023 study by the Federation of European Laboratory Animal Science Associations included responses from ninety-seven animal facilities, and reported just 3,830 animals being rehomed in 2017-2019 [18]. Only 437 rats were rehomed in that period, an average of <2 per facility per year. Perhaps this is because mice and rats are not necessarily widely perceived as potential pets [17,19,20]. However, despite the fact that concerns have been voiced and explored around keeping rats as pets [21,22], evidence suggests that rehomed rats can readily adapt to a new environment [4,5,8,23,24].

In 2023, a rehoming initiative was established at the University of Edinburgh by three of the authors (CQ, IV-F and JM). This led to the formation of a working group tasked to develop policies for rehoming laboratory rats in line with UK government advice and other rehoming initiatives [13]. Intuitively, rehoming seems universally beneficial: the animals live on in a caring environment, and the workers who previously cared for and about the animal may prefer to see the animal rehomed rather than killed (particularly if it was they themselves who would have done the killing). To date, research on rehoming has focused chiefly on guidance, policies, protocols and outcomes in new and well-established rehoming initiatives, or on an individual’s experiences of interactions with rehoming processes and rehomed animals [4–6,8,15,18,23– 27]. These are important research questions, but there has been relatively little research into stakeholders’ perceptions of laboratory rodent rehoming [17,23], and previous research has tended to focus on perceptions of animal facility staff. We wanted to broaden this to explore the views of three key stakeholders: the people who volunteer to rehome these animals, the researchers who normally determine the nature of, and often carry out, the regulated scientific procedures with these animals, as well as the animal technicians and technologists who care for these animals (and who, arguably, know the animals best before they leave the research environment [28,29]). To do this, we carried out semi-structured interviews with stakeholders and subjected interview transcripts to a thematic analysis. We anticipated that by taking a semi-structured approach with a range of stakeholders we would obtain insights that could help inform our working group’s current and future processes and indicate avenues for future reflection and research for our and other rehoming initiatives.

## Materials and methods

The study was approved by the University of Edinburgh Social Research Ethics Group (reference: 2324 SREG 005). Limitations of the study are given in the Supplementary Text.

Semi-structured interviews were conducted by one of the authors (GG) with individuals belonging to one of three groups: people who rehomed laboratory rats from University of Edinburgh facilities (“Rehomers”), animal technicians and technologists based in University of Edinburgh animal facilities (“Technicians”), and PhD students or principal investigators based at the University of Edinburgh (“Researchers”).

All participants were recruited via direct email contact from one of the authors (JM or IV-F). Twelve participants took part in the study: four participants in each of the three groups. Participants gave informed consent prior to participation. Interviews were recorded and transcribed using Microsoft Teams. The interview structure is given in the Supplementary Text. Pseudonymised transcripts were reviewed against the original audio recording on the day of the interview and corrected if necessary. Transcripts were subjected to a thematic analysis using Braun and Clarke’s six-phase method [30]. First, we read and reread the transcripts to familiarise ourselves with their content. Next, each individual transcript was coded to identify features of the data that were relevant to the broad research question, allowing the generation of initial codes. Coding allows the systematic categorisation of segments of data (i.e., interviewee comments) that capture specific concepts. Next, we used Nvivo (Lumivero, Denver, CO, USA) to analyse codes, grouping related codes into themes. These themes are higher-order patterns or concepts derived from the codes and representing recurring ideas and topics relevant to the research question. Themes provide a broad interpretation of the data by grouping related codes together, representing underlying meaning of the data and offering insights into the research question. All codes from the initial coding phase were classified as representing either a supportive or a cautious perception or attitude and were incorporated into a theme. It is important to note that ‘cautious’ codes often referred to challenges around rehoming rather than expressing opposition to it. Next, we reviewed themes, combining similar themes if necessary, ensuring that each theme was distinctive and coherent in terms of its content. To explore quantitative differences between interviewee groups’ codes, we quantified the total number of supportive and cautious codes in each interviewee group, then ranked the codes by frequency in each theme (Supplementary Data). In line with requirements to minimise the risk of deanonymisation, all participant quotes given below are paraphrased.

## Results

We documented a range of codes that we allocated to a total of seven supportive themes and eleven cautious themes. We then grouped these themes into three families: Animals’ interests, People’s interests, and Institutional interests (Table 1) and quantified the total number and category of codes expressed by each interviewee group for each theme (Figure 1).

**Table 1.** Themes on perceptions of rehoming laboratory animals.

| Theme family | Supportive themes | Cautious themes |
| --- | --- | --- |
| Interests of the animal | Animals can 'retire' instead of being killed. | Issues with rehomer selection. |
|  | Quality of life may be higher when rehomed compared to in an animal facility. | Issues around the animal's welfare after rehoming. |
|  | Interactions with institution staff socialises laboratory animals. | Issues around the animal's adaptability to the rehoming environment. |
|  |  | Rehoming is not possible because of the animal's genotype/phenotype, or how it was used in scientific studies. |
| Interests of the people involved | Rehoming is a straightforward process. | Negative feelings associated with not rehoming animals. |
|  | Positive emotions around the act of rehoming. | Increased rehoming-related workload for staff. |
|  |  | Concern around rehomers' health. |
|  |  | Concerns around the affordability of rehoming for rehomers. |
| Interests of the institution | Potential for increased transparency between the institution and the public. | Rehoming draws attention to the use of animals in research. |
|  |  | The limited capacity to rehome. |
|  |  | The option to rehome is not widely advertised. |

**Figure 1.**
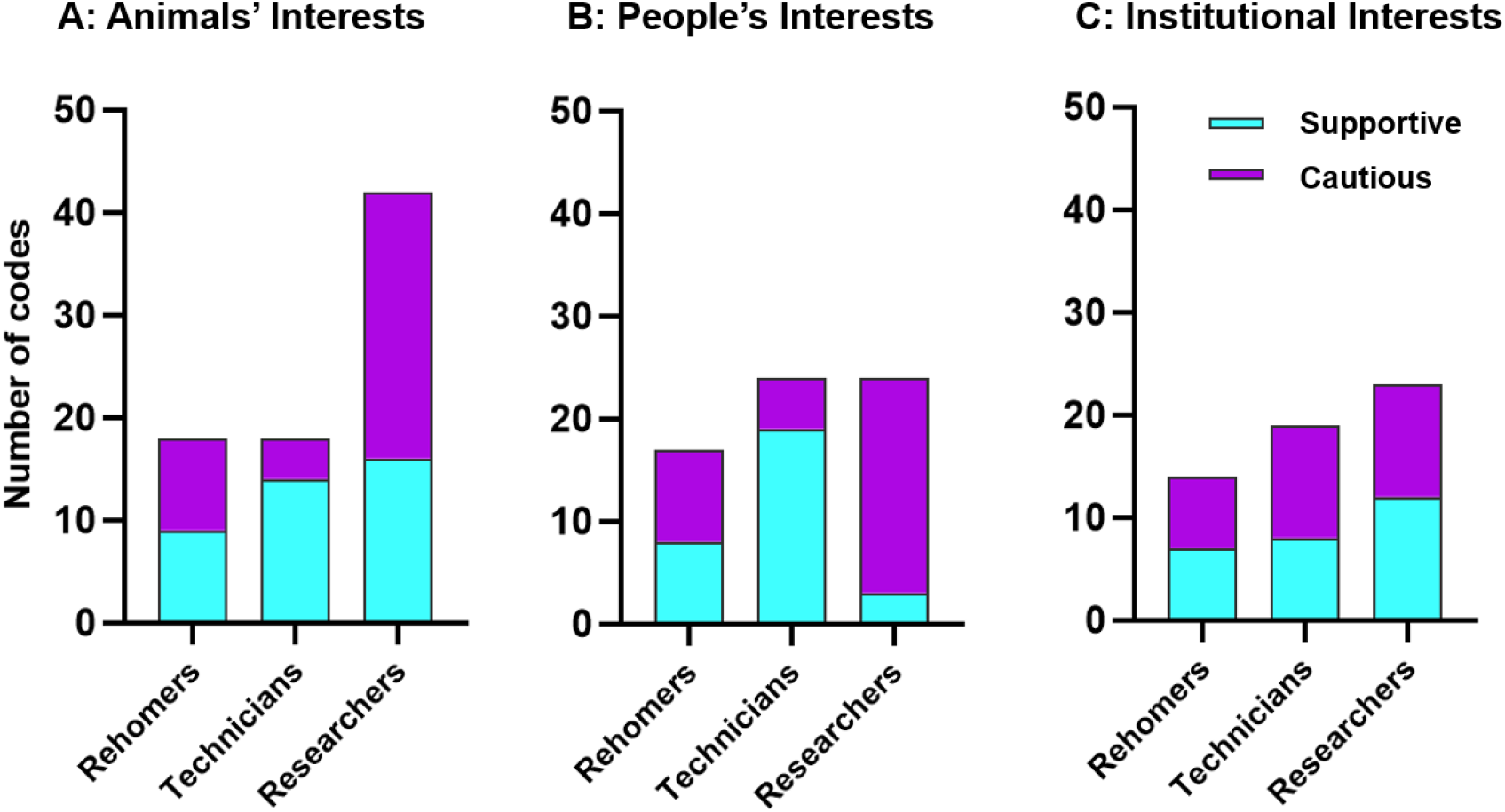
Total numbers of supportive and cautious codes documented for each interviewee group in (A) the Animals’ Interests family, (B) the People’s Interests family, (C) the Institutional Interests family.

All groups expressed both supportive and cautious codes across all families of themes. In the Animals’ Interests family, Technicians primarily expressed supportive codes. Researchers expressed more than double the number of codes compared to the other two groups, and cautious codes dominated. Rehomers expressed an equal number of supportive and cautious codes. In the People’s Interests family, a similar pattern was seen, with Technicians predominantly expressing supportive codes, Researchers predominantly expressing cautious codes, and Rehomers expressed a roughly equal number of supportive and cautious codes. In the Institutional Interests family, all three stakeholder groups expressed a roughly equal proportion of supportive and cautious codes, with the Technicians expressing the largest proportion of cautious codes.

To explore the priorities of each group, we next documented which specific codes were most frequently expressed by each group (Supplementary Data). In terms of Animals’ Interests, all three groups frequently commented regretfully on the usual practice of killing animals at the end of a study, and on the potential benefits of rehoming on the welfare of these animals. One interviewee reflected on having to kill their animals after the study:

> *That’s no one’s favourite day at work. You come into work, and you think: oh my god, I have to do that today. [Researcher]*

Another interviewee, reflecting on the potential for rehoming, said:

> *I was trained by an amazing person who always used to tell me that you have a choice to work here.*
>
> *[The animals] don’t; they have no choice. So why not just let them go, to be comfortable somewhere else instead of being here? [Technician]*

Interviewees also raised the normal limitations of the facility environment, and compared with the level of care available in a rehoming environment:

> *We’d be caring for several animals in a room, instead of one-to-one. They’d get more taken care of in the home. [Technician]*
>
> *They get varied food, better exercise and handled a lot more than they do in the labs. [Technician]*

The Rehomer group commented widely on the rehomed animal’s welfare, discussing rehoming as a positive alternative to death. For example, one talked of:

> “*the little creatures that you’ve taken care of, knowing that they will go from their study to go on to live a full, happy life rather than being euthanised at the end of the study”. [Rehomer]*

In terms of People’s Interests, the Technician and Rehomer groups frequently expressed codes related to their positive emotions around the act of rehoming, and negative emotions about animals that could not be rehomed. Interviewees in the Technician group made several comments on the direct and positive impact of rehoming on them:

> *For me personally, it just makes me smile, you know, and I’ve done some good. Because this can be a hard job, psychologically and mentally it can be quite difficult at times. It’s a big pressure off my mind knowing I don’t have to take them down that route [euthanasia], it’s very beneficial to me. [Technician]*
>
> *[Rats are] amazing pets so the fact that you can let them live a happy life with somebody else, that is a great, great idea. And it prevents us from killing them, it’s amazing. [Technician]*

One of the Technicians talked of their delight around one of the rehoming events:

> *Yeah, that day! It’s quite an exciting thing whenever we’re getting some that are getting rehomed, that means we get to go through and think: ok, who’s going to get rehomed, who’s able to go? [Technician]*

Interviewees in the Researcher group also acknowledged the positive impact on themselves. For example, one commented:

> *It feels good as a researcher to be doing something: that retirement stage after their service, and you see where they’re going to live, hearing back from them how they’ve settled in, and it’s just very nice. I would have loads, I’d be taking them home every week. [Researcher]*

A Rehomer commented on the impact on the animals’ former carers when sharing updates of rehomed animals’ progress:

> *Occasionally I’ll send photos and things just to show them, some of the technical staff have said that they’ve been really touched when they saw those photos of them just having fun and doing ratty things. [Rehomer]*

This was reflected in one of the Technician’s comments:

> *It’s nice to get to see them in their new environment. I’d say they look a bit happier; they definitely look as though they’ve got more going on. [Technician]*

However, the Researcher group frequently expressed concerns about the pragmatic aspects of rehoming and its governance. For example, the workload involved, capacity issues, how rehomers are selected, potential impacts on rehomers’ health, and post-rehoming animal welfare challenges. For example:

> *Someone that doesn’t understand the responsibility of rehoming, which is always a concern, what if they don’t know they need to be kept in social groups or have other pets which is stressful. [Researcher]*.

In contrast, the Rehomer group raised the pragmatic aspects of rehoming much less frequently.

In terms of Institutional Interests, interviewees from all stakeholder groups commented on the possible benefits of a rehoming programme to openness, recognising its potential as a policy and practice that could demonstrate a commitment to welfare:

> *Any use of animals in science is always a little controversial for the public. So, I think for the University, anything that shows that they’re doing what they can to reduce harm to animals, to give them opportunities to live out their lives, I think is a good thing. [Rehomer]*
>
> *It’s good for the University’s reputation, it shows the university is engaging with, and more involved with, the animal welfare side of things. [Technician]*
>
> *I think it’d be a nice public engagement exercise. Showing that when animals are finished their experiments, they could go on to be family pets. [Researcher]*

But interviewees also recognised potential risks. Particularly the risk that rehoming will draw attention to the fact that animals are used in research:

> *It does remind people that laboratory rats are being used in the University, when people don’t generally think they are. [Rehomer]*
>
> *If we are trying to be positive: look, we’re rehoming this number of animals. Well, what happened to them before? Why are you using them? [Researcher]*

One Rehomer raised the issue of ‘waste’ [12], and questioned whether a rehoming programme might be perceived as a dubious way to mitigate for it if:

> “… *it was seen in terms of people thinking they have more rats than they need and they’re just giving them out to whoever says they want them” [Rehomer]*

Another interviewee alluded to the potential risks to people who rehome laboratory animals:

> *It’s not really good to hide and pretend … but I can see why some people might be a little bit fearful if you’ve rehomed a lab animal and people find out*” *[Researcher]*

## Discussion

We carried out semi-structured interviews to explore perceptions around laboratory rodent rehoming with three groups of stakeholders: Rehomers, Technicians and Researchers. A thematic analysis demonstrated clear support across all three groups for laboratory rodent rehoming, though different stakeholder groups had different foci of attention. Rehomers and Technicians tended to focus on the benefits to the animal and the institution. Researchers shared this focus too, but these comments were often expressed alongside concerns around the welfare of the rehomed animals, rehomers’ well-being, the fate of animals that cannot be rehomed, and the potential impacts of a rehoming scheme on institutional reputation. We discuss the Animals’ Interests, People’s Interests, and Institutional Interests families of themes further, mapping them onto (1) approaches to laboratory animal ethics and welfare, (2) manifesting a Culture of Care, and (3) opportunities around institutional openness.

First, we consider our findings in the context of ethics and welfare. In the introduction, we asserted that being rehomed is a better outcome for the animal than being killed. This is in line with, for example, Martha Nussbaum’s view that death is a harm for an animal if that animal is aware of and values the “temporal unfolding” of their life [9]. There is emerging evidence that rats have complex cognitive capabilities, including a capability to re-experience past events and imagine future events [31] and a capability to plan future behaviour [32]. Rats appear to value their own existence by, for example, seeking what we might imagine as pleasurable experiences and avoiding harmful experiences [33–35]. Accordingly, it seems at least possible that death is a harm for rats. However, and perhaps surprisingly, the humane killing of laboratory animals is not regarded by some as a welfare issue [36]. For example, Broom describes killing as “an important ethical question but it is not an animal welfare issue. The animal welfare issue is what happens before death” [37]. Similarly, Franco discusses (but does not support) the view that “painless killing of laboratory animals poses no ethical problems, as nonexistence implies absence of negative experiences” [38]. Rollins suggests that simply by virtue of being a laboratory animal, “[the animal’s] *telos* is regularly violated, hence it is not happy and painless death is no harm to it” [39]. Interestingly, perhaps this implies that death would become a harm if that animal’s identity changed – as it arguably does in the act of rehoming – from ‘laboratory animal’ to ‘companion animal’. At any rate, ending an animal’s existence entails both the loss of future opportunities to express its capabilities, and the absence of future positive experiences. If euthanising in order to prevent suffering *is* considered as a welfare issue, it is difficult to understand why the opposite - prolonging the life of an animal that is not experiencing suffering and has the potential to have positive experiences - is not. One of our interviewees recognised this, and – imagining addressing their animals at the end of a study – raised the quandary of rehoming some animals and killing others:

> *Some animals have a nice happy retirement and the rest of you, I’m really sorry, but I’m going to have to kill you. It’s very questionable, isn’t it? [Researcher]*

The view that it *is* ethically correct to prolong animals’ lives has been expressed fairly widely [9,23,40–44], provided, of course, that these animals experience a good level of welfare. We contend that rehoming, when a ‘good’ rehoming is possible, is preferable to the death of the animal, and that this creates a moral incentive to provide opportunities for rehoming.

But the potential benefits of rehoming go beyond a simple extension of lifespan. There are limitations to the amount and type of care that can be provided in an animal facility environment. Though these are often rooted in limited resources rather than a lack of will or motivation to provide care; bonding between animal technicians and laboratory animals and the high level of care they provide is well-documented [23,45,46]. Rehoming can give the opportunity to prolong life in a larger, more enriched environment that can benefit the social and physical well-being of the rats, with opportunities for rats to experience new activities, foods and social interactions [47]. This is in line with broader emerging ideas around promoting positive welfare and giving animals in our care ‘a life worth living’ [48,49], rather than the traditional approach in the animal research context of seeking only to minimise suffering.

There are challenges around providing a ‘good’ rehoming too, of course. For example, one contributor in a different study of rehoming [23] expressed their perception that standards of care in animal facilities are, at least, consistent: “*I think a companion animal has a bit more of a lottery in their existence perhaps than these research animals*”. Arguably, being a laboratory rat is also lottery; a given rat’s ‘job’ could be confined to breeding or taking part in a behavioural study involving rewarding interactions with humans and other rats. On the other hand, the ‘job’ could involve recovery surgery and potentially severe procedures. Nonetheless, rehomed animals *are* being brought into a new environment, and differences between animal facility environments and routines and those of the rehoming environment require the animal to adjust. This transition may be difficult for some rats, and the adjustment to the new environment stressful. Rats may be provided with more care, attention and resources from their new owner post-rehoming, but concerns persist that animals’ experiences may be negatively affected by the less regulated standard of care given in private homes compared to the laboratory, particularly with novice rehomers. Also, simply because rehomed animals will likely live to an older age, they will have an increased chance of aged-related health issues that diminish their quality of life. However, one Rehomer in our study gave a very positive example of how well the animals adapted to their new environment and house-mate:

> “*The rats weren’t phased by anything, the hoover, my pet cat, they took it all in their stride*”. *[Rehomer]*

All interviewee groups raised the issue of capacity, questioning how many rats can realistically be rehomed. For example:

> *There’s an element of guilt. You can’t do it yourself, and there’s only so much time you can give to finding these animals a home. And if I could take them myself, I would be overrun with rats. [Researcher]*

There are only around 100,000 rats kept as companion animals in the UK [21] (compared to an estimated 21 million cats and dogs [50]). This implies there is a very limited capacity for rehoming rats. It is not clear how to address this capacity issue, but it may be interesting to explore how different institutions approach the recruiting of rehomers. Significant efforts have been made to provide frameworks for effective rehoming [15], but further research exploring concerns around opening rehoming initiatives to people outside the institution, particularly the ability to evaluate potential rehomers when there is no one from the institution to vouch for them, may be of use to support the development of internal institutional policies on rehoming.

This capacity issue has led some researchers (and one of our interviewees) to suggest alternative ethical uses of laboratory rodents, for example, using humanely-killed laboratory rodents in place of rodents bred solely to be used as food for pet or captive reptiles and birds [51]. However, this proposal raises several ethical issues in itself, not least to the facility supplying these animals [8], and whether the keeping of ‘exotic’ (or, indeed, any) animals as pets or in captivity should be supported [19,52], and, if it is, whether it matters to that animal whether its prey is alive or dead when it is presented to it [53].

It is important that rehoming does in fact benefit the animal, especially when we benefit from the research they are involved in, and they do not [4,17]. However, rehoming may also benefit the people who work with these animals; potentially in diverse ways, from partially alleviating compassion fatigue [4,46,54] to accumulating political or social capital in wider society [4,5]. Accordingly, we next consider our findings in terms of manifesting a Culture of Care.

Culture of Care has been defined as “an establishment-wide commitment to improving animal welfare, scientific quality, care of the staff, and transparency for all stakeholders, including the public” [55]. Here we focus on the impact of rehoming on some of the staff who are responsible for many of the interactions with laboratory rodents. In our study, Technicians rarely raised challenges related to the workload involved in rehoming, despite them being responsible for a significant amount of this work. This reminds us of the important role that animal technicians have in helping understand which specific, individual animals should be rehomed: as Skidmore points out “The formation of a relationship [with these animals] in turn makes staff more likely to instigate the necessary procedures, such as training and socialisation, to prepare animals for rehoming” [17].

As well as having major focus on the benefits of rehoming to the animals and to institution openness, this stakeholder group also remarked in positive terms on the impact rehoming had on them. Beth Greenhough and Emma Roe noted that “at the heart of the role of [animal technicians] is the tension between the need to provide care and how this is harnessed by laboratory animal science for utilitarian ends through the endeavour of producing compliant and useful bodies for science” [29]. Part of this tension are the acts of caring for then killing animals at the end of a study, responsibilities that are often associated with stress and compassion fatigue [46,54]. Greenhough and Roe have highlighted the stress felt by technicians when asked to kill healthy animals: “what comes across strongly from [technicians’] comments is the value they place on care to support life as opposed to valuing their capacity to end life” [28], echoed by Gail Davies’s ideas of “shared suffering” and its effects on human and animal well-being [56]. Because many aspects of technicians’ roles are very challenging, they may see rehoming as one activity with a very clear positive and rewarding outcome for them as well as for the animals. For example, one of our interviewees in the Technician group commented:

> “*The rats are going to have a much better experience. I’ve seen the cages that they were going to be rehomed into, and it all looks just amazing. They’re going to be running about, climbing, burrowing, nesting, that they can’t really do in [laboratory] cages*”. *[Technician]*

Rehoming, as an alternative to killing, may be valuable to technicians, encouraging a deeper engagement with a Culture of Care. This is in line with Greenhough and Roe’s observation that technicians can perceive “animal lives in ways that bear witness not only to animal suffering, but to animal agency … hanging on to the speculative desire, if not the immediate possibility, of seeking other ways of living together” [29]. Perhaps rehoming is one of these “other ways”.

In our study, Rehomers did not raise any concerns around impacts on their own health, well-being or their willingness to rehome. This may be because potential rehomers self-select in terms of their ability and motivation to bring these animals into their home and care for them. However, others have noted challenges associated with Rehomers’ expectations, the costs involved, and the efforts needed to maintain animal health and socialisation [15] and the morality of rehoming [8]; any rehoming scheme should seek to support rehomers in those respects. Unexpectedly, the Researchers in our study frequently expressed concerns about the pragmatic aspects of rehoming. This surprised us because most of the practical and administrative workload involved in rehoming does not fall on this group. It may be the case that are some interviewees in the Researcher group were not fully aware of what the rehoming programme involves, for example, in terms of evaluation of rehomer suitability.

We next turn to the idea of publicising rehoming as part of an institutional openness policy. Institutional openness is normally defined as the sharing of information in the public domain on how and why an institution uses animals in scientific research. Our interviewees highlighted a tension between a desire for openness and a desire to avoid controversy. This is understandable given the most recent polling in the UK on perceptions around animal research [57]. Around two-thirds of respondents accepted the use of animals, but there was also significant concern for animal welfare in the research context: 38% said that animals should not be used in any research on animal welfare grounds. When asked about the organisations involved in animal research, the primary characteristic attributed to these organisations was “secrecy”, and only a quarter of all respondents believed that these organisations are well-regulated. Although caution is needed around using public opinion polls to legitimise or give authority to any side of arguments around using animals in research [58], against this background it seems clear that institutions need to navigate a careful balance between openness, protecting their workers, the risks of alienating the public and the anticipated consequent reduction in public support for using animals in research. This balance was discussed by Pru Hobson-West and Gail Davies as an effort to “reduce *societal* pain, suffering, distress, and lasting harm potentially caused by laboratory animal science”: an interesting and innovative parallel to the way application of the 3Rs seeks to minimise pain, suffering and distress to animals [59].

Other research has noted that challenges around openness are recognised by animal facility workers. In one study of facility managers at a Canadian institute [60], many participants recognised that a lack of openness can give an impression of secrecy, and almost all participants expressed a wish to be transparent, and that “they wanted to feel like they had nothing to hide”. Interestingly, and perhaps overly optimistically, this desire was often motivated by a belief that transparency would increase public scientific literacy, and that this, in turn, would result in wider support for the use of animals. However, this group also expressed concerns around their own safety and security if openness was increased. A related study of vets working in Canadian universities’ animal facilities [61] also expressed the belief that openness may lead to increased public support, but the authors described other, more nuanced, views on openness, including a clear recognition that openness could result in negative impacts on individuals and the institution, and that a strategic and selective form of transparency they coined “agenda-driven transparency” might be used to positively influence public support. Two studies of UK vets [62,63] reflected many of the findings from these Canadian studies, but also highlighted specific concerns around the consequences of openness, particularly towards the common representation of vets as the advocate, or even guarantor, of animal well-being, and how this perception might be affected if the complexities around their roles in animal research became part of a wider discourse.

Our study supports the idea that those directly affected by increased openness have mixed feelings about its benefits. While acknowledging that this article is in itself a form of (limited, peer-oriented) openness, we next ask: should laboratory animal rehoming programmes be publicised? This question was explored extensively by Tess Skidmore [8]. In line with our study, she found that rehoming can enhance a culture of care and describes her research participants’ “frustration and anger when rehoming is not attempted for fear of reputational damage”, and that deciding not to rehome animals diminishes “the circulation of hope, compassion and care”. However, she also reports that rehoming activities “can also be the catalyst for cultures of anxiety and risk”, describing how publicity around rehoming can draw unwanted attention to practices in animal research (including the practice of rehoming itself) and to the people who carry out these practices, particularly if these animals are described as “traumatised” or “rescued” from a laboratory. Interestingly, she explores the idea that rehoming “thrives through a culture of discretion rather than openness”, and that, in contrast to the conventional claims that openness enhances trust between groups with different interests, she suggests that trust in the context of rehoming may be “promoted through the security and assurance instilled by a lack of openness” (this applies to the public’s trust of institutions, and to institutions’ trust of the public). Accordingly, she concludes – and we agree – that there is significant potential for publicity being a deterrent to institutions introducing rehoming activities.

In terms of future research on rehoming, animal technicians have been identified as a group with useful perspectives and understanding of the human-laboratory animal relationship [29], and future initiatives co-created with this group may reveal how these relationships can feed into rehoming processes to benefit all stakeholders. More specifically, stakeholders’ pragmatic concerns about rehomed animals’ well-being could be addressed by collecting data and anecdotes from rehomers, potentially sharing these in the manner of Eva Meijer’s work on living with rehomed laboratory mice; on how they live, their individualised behaviours and personalities, and their interactions with her and each other [24]. These anecdotes could also be used to inform and enhance approaches in institutional animal facilities that seek to recognise the full capabilities of animals and promote positive welfare.

However, these approaches highlight a tension: by offering laboratory rats as subjects for rehoming, we are implicitly suggesting that they make good companions. Why do they make good companions? Because they are social, intelligent, affectionate and playful. So, thinking about *why* rats may ‘deserve’ to be rehomed may lead to a shift in perception of what attributes and capabilities they possess. This shift could, in turn, make more explicit the difficulties in reconciling an appreciation of these attributes and capabilities with some of the ways in which these animals are used in research. Exploring and managing this tension may be one of the main challenges that arises from rehoming.

## Acknowledgements

The authors thank all study participants, members of the University of Edinburgh 3Rs and Culture of Care Committee and members of the University of Edinburgh Lab Rodent Rehoming Working Group. JM thanks Nairn Menzies for discussions that stimulated the development of the rehoming initiative. For the purpose of open access, the authors have applied a Creative Commons Attribution (CC BY) licence to any Author Accepted Manuscript version arising from this submission.

