## Supplementary text for "Rehoming laboratory rats: Exploring perceptions of rehomers, animal technicians and biomedical researchers"

### 1. Plans for semi-structured interviews

*1.1 Plan for Researcher and Technician participants*

1. Welcome, obtain consent, participant and facilitator introductions, briefly describe plan for the discussion, including clarification of what we are referring to by ‘rehoming’. (~5 min)

2. What do you see as the positive aspects of rehoming? (~20 min)

If not already discussed, raise:

- Potential benefits to animal.
- Potential benefits to people involved in study and/or care.
- Potential benefits to institution.

3. What do you see as the negative aspects of rehoming? (~20 min)

If not already discussed, raise:

- Potential impacts on animal.
- Potential impacts on people involved in study and/or care.
- Potential impacts on institution.

*1.2 Plan for Rehomer participants*

1. Welcome, obtain consent, participant and facilitator introductions, briefly describe plan for the discussion, including clarification of what we are referring to by ‘rehoming’. (~5 min)

2. What do you see as the positive aspects of rehoming? (~20 min)

If not already discussed, raise:

- Potential benefits to animal.
- Potential benefits to person rehoming.
- Potential benefits to people involved in study and/or care.
- Potential benefits to institution.

3. What do you see as the negative aspects of rehoming? (~20 min)

If not already discussed, raise:

- Potential impacts on animal.
- Potential impacts on person rehoming.
- Potential impacts on people involved in study and/or care.
- Potential impacts on institution.

**2. Limitations**

This study took place as part of one of the author's (GG) final year research project in her undergraduate degree programme. This context constrained the number of interviews that could take place. Despite this limitation, we documented a range and diversity of codes between individuals and groups. However, compared to data obtained from a larger sample, our data may be relatively heterogeneous.

Our participant sample was not randomly selected; we directly approached colleagues who we believed had an interest in issues around laboratory animal science and welfare. However, interviews took place independently of each other, and as far as we are aware, participants were not aware of other interviewees' identities and did not interact with each other in ways that might influence how they engaged with our study.

Issues around demand characteristics (for example, the apprehensive participant role [1]) are difficult to identify and mitigate [2], but the interviewer sought to find a balance between encouraging interviewees to express themselves while trying to avoid expression of her own thoughts, beliefs and values in a way that might influence the interviewee. However, an interviewee who was indifferent, or even opposed, to rehoming may not have felt comfortable expressing such views during the interview.

The interviewer's identity may also be a factor in how interviewees engaged in our study. These concerns are similar to those raised by Tess Skidmore in her PhD thesis on rehoming [3]: that she "*felt very aware of my younger age and less advanced career stage whilst conducting the interviews*". Skidmore also reported that that she may have been perceived as "*an outsider in that I am young, do not have a background in biological sciences and do not actively work in the field of animal research*", and that interviewees may have had "*the perception that I did not agree with animal research given my interest in rehoming*". The interviewer in our study was relatively young and at an early career stage but did have a background in biomedical sciences.
